# Gene transfer of human cardiomyopathy β-MyHC mutant R403Q directly alters intact cardiac myocyte calcium homeostasis and causes hyper-contractility

**DOI:** 10.1101/2024.07.31.605903

**Authors:** Todd J. Herron, Eric Devaney, Guadalupe Guerrero-Serna, Lakshmi Mundada, Joseph M. Metzger

**Author notes:** Correspondence to Todd J. Herron, PhD, Cardiovascular Regeneration Core Laboratory, 2800 Plymouth Road, B26-223N, Ann Arbor, MI USA.

## Abstract

The R403Q mutation of human cardiac β-myosin heavy chain was the first missense mutation of a sarcomeric protein identified as being causal for hypertrophic cardiomyopathy (HCM), in humans. The direct effect of the R403Q mutant myosin on intracellular calcium homeostasis and contractility is not fully known. Here we have used *in vitro* gene transfer of the R403Q mutant human β-myosin to study its direct effects on single intact adult cardiac myocyte contractility and calcium homeostasis. In the first experiments, adult cardiac myocytes transduced with the R403Q mutant myosin recombinant viral vectors were compared to myocytes transduced with wild-type human β-myosin (wtMYH7). Efficiency of gene transfer was high in both groups (>98%) and the degree of stoichiometric myofilament incorporation of either the mutant or normal myosin was comparable at ∼40% in quiescent myocytes in primary culture. Sarcomere structure and cellular morphology were unaffected by R403Q myosin expression and myofilament incorporation. Functionally, in electrically paced cardiac myocytes, the R403Q mutant myosin caused a significant increase in intracellular calcium concentration and myocyte hyper-contractility. At the sub-cellular myofilament level, the mutant myosin increased the calcium sensitivity of steady state isometric tension development and increased isometric cross-bridge cycling kinetics. R403Q myocytes became arrhythmic after β-adrenergic stimulation with spontaneous calcium transients and contractions in between electrical stimuli. These results indicate that human R403Q mutant myosin directly alters myofilament function and intracellular calcium cycling. Elevated calcium levels may provide a trigger for the ensuing hypertrophy and susceptibility to arrhythmia that are characteristic of HCM.

## Introduction

Hypertrophic cardiomyopathy (HCM) is an inherited autosomal dominant disease characterized by unexplained cardiac hypertrophy and myocyte and sarcomere disarray ^1,2^. HCM is the most common cause of exercise-related sudden death in young athletes ^3^. Molecular genetic linkage analyses in large families have demonstrated that HCM is a disease of the cardiac sarcomere ^4–6^. The most affected sarcomere genes are among those that make up the thick filament of the sarcomere, namely *MYBPC3* (cardiac myosin binding protein-C) and *MYH7* (β-myosin heavy chain, β-MyHC) ^7^. β-MyHC is a molecular motor isoform of cardiac muscle that drives myocardial contraction ^8,9^. The R403Q missense mutation of the human β-MyHC molecule was the first point mutation of a sarcomere protein reported to cause HCM ^6^. This mutation is associated with a poor prognosis and high incidence of cardiac arrhythmias and sudden cardiac death ^10,11^. The underlying mechanisms whereby this point mutation of myosin leads to arrhythmia and sudden cardiac death, however, is not completely understood.

Since the discovery of the R403Q myosin mutation much effort has been devoted to determining how this mutant directly affects myosin motor function and how this might, in turn, alter intracellular calcium flux to trigger arrhythmia. A full understanding of the R403Q mutant myosin effect on motor protein function remains incomplete. Studies have suggested that myosin motor function is impaired ^12–16^, while others suggest a gain-of-function for the myosin motor ^17–21^. The effect of this mutant myosin on intracellular calcium homeostasis is also unclear. One study using a transgenic mouse model demonstrated that calcium levels were elevated in electrically paced trabeculae isolated from mutant mice ^22^. In contrast, there are reports that cardiac myocyte intracellular calcium levels are lower in these same mutant mice ^23,24^. One key limitation of the transgenic mouse model is that the disease-causing mutation was expressed in the functionally and structurally distinct α-MyHC motor isoform ^8,25,26^, while the human disease-causing mutation is expressed in the β-MyHC motor isoform. Indeed, an examination of this mutant myosin reported that the functional effects of the R403Q mutant are different in the α- vs. β-MyHC backbone ^27^. Further, in transgenic animal models it may be difficult to distinguish primary effects caused by the R403Q mutant myosin from secondary compensatory changes that may occur throughout the lifespan of the animals ^28^. Thus, a key question remains as to whether there are direct effects of the R403Q mutant myosin in the context of the human β-MyHC molecule when incorporated into intact myocytes.

Our aim here was to examine the acute effects of the human R403Q mutant β-MyHC molecule on adult cardiac myocyte intracellular calcium homeostasis and contractile function by using *in vitro* gene transfer. We generated recombinant adenoviruses to express the full-length wild type human β-MyHC gene or the HCM-causing R403Q mutant myosin in adult cardiac myocytes *in vitro*. Strength of the *in vitro* gene transfer approach employed here is that the cardiac myocytes are quiescent and contractile protein expression is stable ^29–31^ during adenoviral-based transgenesis of the mutant myosin molecule. We report here that the R403Q mutant myosin increases intracellular calcium concentrations, increases myofilament calcium sensitivity, increases spontaneous calcium release from the sarcoplasmic reticulum and causes hyper-contractility in adult rodent cardiac myocytes.

## Methods

### Mutagenesis and Recombinant Adenovirus Production

Details for the cloning and production of adenovirus to express the full length human β-MyHC motor protein have been reported previously ^31^. The mutated amplimer sequence for the R403Q missense mutation in the context of the human *MYH7* gene was identified from the CardioGenomics database (Genomics of Cardiovascular Development, Adaptation, and Remodeling. NHLBI Program for Genomic Applications, Harvard Medical School. URL: http://www.cardiogenomics.org). The mutation is in exon 13 and the codon change is CGG>CAG. Site directed mutagenesis was performed by polymerase chain reaction using protocols of the Stratagene Quik Change mutagenesis kit to introduce the R403Q mutation into the human MYH7 gene. Successful site directed mutagenesis was confirmed by DNA sequencing of the full-length cDNA (University of Michigan DNA sequencing core) and translation of the wild-type and mutant myosin are shown in Figure 1. Homology is highlighted in yellow, the missense mutation is apparent at amino acid position 403 (white). The AdMax system (Microbix) was used to generate recombinant adenoviral vectors as previously described ^31,32^.

**Figure 1.**
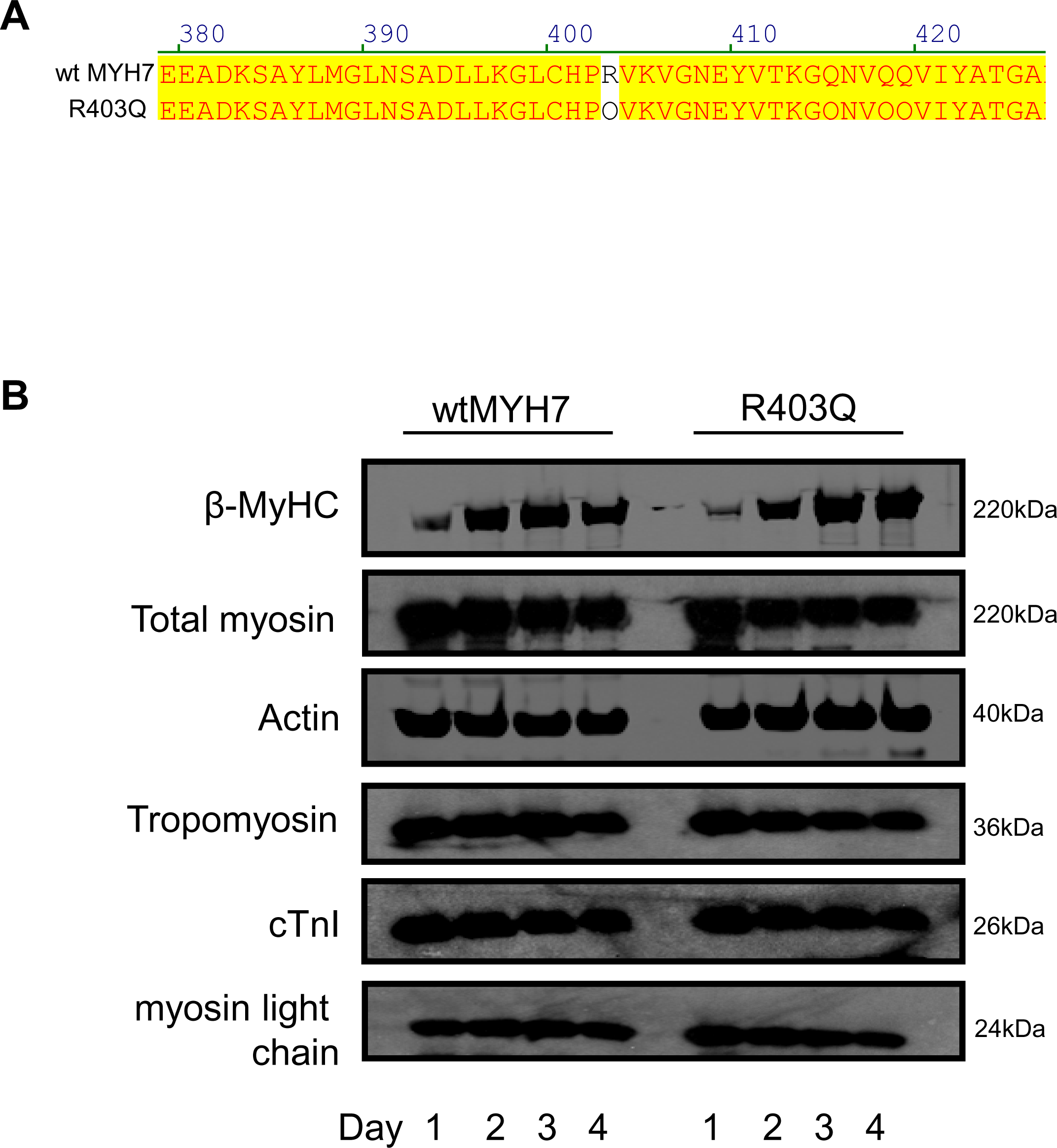
R403Q mutant human myosin expression in adult cardiac myocytes. **(A)** Site directed mutagenesis of the human MYH7 gene was confirmed by DNA sequencing and translation of the expression cassette driven by the CMV promoter is presented. Homology with the wild type (wtMYH7) sequence is highlighted in yellow and R to Q mutation shown (clear text). **(B)** Western blotting shows that both R403Q and wtMYH7 β-MyHC protein expression increased over time in culture, with no significant change in total myosin content. The expression of other key myofilament proteins were unaltered by *MHY7* gene transfer. Myosin light chain is MLC2V (RLC).

### Adult cardiac myocyte isolation and primary culture

Adult rat cardiac myocytes were isolated from female Sprague Dawley (Harlan, 150-200g) rats and cultured and gene transfer was performed as reported previously ^30,31^ with minor modification. To increase myocyte viability, blebbistatin (25µmol/L, Toronto Research Chemicals, Inc.) was added to the calcium free collagenase solution and perfused through the heart during enzymatic digestion. This specific myosin II inhibitor was washed out after cell isolation. Blebbistatin has been shown to enhance cardiac myocyte viability in primary culture without affecting adenoviral gene transfer ^33^. Calcium tolerant cardiac myocytes were plated on laminin coated coverslips (∼2×10^4^ myocytes/coverslip) and treated with R403Q or wtMYH7 adenovirus in M199 media. Multiplicity of infection (MOI) was 500 for each virus. Myocytes were maintained in primary culture in M199 media for up to four days. All animal use agreed with the guidelines of the Internal Review Board of the University of Michigan Committee on the Use and Care of Animals. Veterinary care was provided by the University of Michigan Unit for Laboratory Animal Medicine.

### Contractility Measurements in Single Cardiac Myocytes

Functional analysis of single cardiac myocytes was performed on day 3 following *in vitro* gene transfer of the wild-type human *MYH7* gene (wt*MYH7*) or the R403Q *MYH7* mutant myosin. We have reported previously using identical experimental conditions that myosin transgene expression represents a stoichiometrically conserved replacement of ∼40% with no changes in the total myosin content by day 3 after adenoviral gene transfer ^31^. Sarcomere shortening and re-lengthening of field stimulated myocytes (50V, 1Hz) was recorded in real time (240Hz) using an IonOptix camera and software. Intracellular calcium transients were measured simultaneously (1kHz) using the ratiometric calcium sensitive probe fura-2AM (2µmol/L). Full details of this system have been reported elsewhere ^30,31^.

For determination of sarcoplasmic reticulum calcium content 20mmol/L caffeine was rapidly applied to myocytes immediately after a bout of steady state pacing at 1Hz. Caffeine was rapidly applied to a myocyte manually using a gravity fed capillary tube perfusion system (Warner Instruments, SF-77B Perfusion Fast-Step) as reported previously ^34,35^. Fractional SR calcium release was calculated by dividing the 1 Hz pacing transient amplitude by the amplitude of calcium released by caffeine application. Responsiveness of myocytes to β-adrenergic stimulation was determined after isoproterenol (10nmol/L) treatment.

Myofilament calcium sensitivity and isometric cross-bridge cycling kinetics were measured using single-membrane permeabilized (1% Triton) myocytes attached to a force transducer and length controller, as previously described ^31^.

### β-MyHC Protein expression

#### Immunofluorescence

*MYH7* gene expression and sarcomere incorporation were probed using a β-MyHC specific antibody (ATCC® number: CRL-2046™, 1:1000) as before ^31^. High resolution confocal imaging was performed using a FluoView™ FV500 laser scanning confocal microscope (Olympus).

#### Western Blotting

Myocytes were harvested in Laemmli sample buffer and proteins were separated by SDS-PAGE. Separated proteins were transferred to nitrocellulose membrane and probed with antibodies specific for β-MyHC or other key myofilament proteins. Total myosin was probed using the MF20 antibody which recognizes both cardiac myosin isoforms. The MF20 monoclonal antibody (1:1000) developed by Donald Fischman was obtained from the Developmental Studies Hybridoma Bank developed under the auspices of the NICHD and maintained by The University of Iowa, Department of Biology, Iowa City, IA 52242. Western blots were also probed with the monoclonal 5C5 actin antibody (Sigma, 1:2500), Tm311 tropomyosin antibody (Sigma, 1:5000), cTnI antibody MAB 1691 (Chemicon, 1:1000) or the cardiac myosin light chain 1 antibody MLM527 (Abcam680, 1:2500). Horseradish peroxidase labeled secondary antibody (BD Pharmingen™, 1:2500) was used in combination with enhanced chemiluminescence for antibody detection. The human β-MyHC R403Q missense mutation did not affect antibody binding to the human β-MyHC protein (Figures 1 and 2).

**Figure 2.**
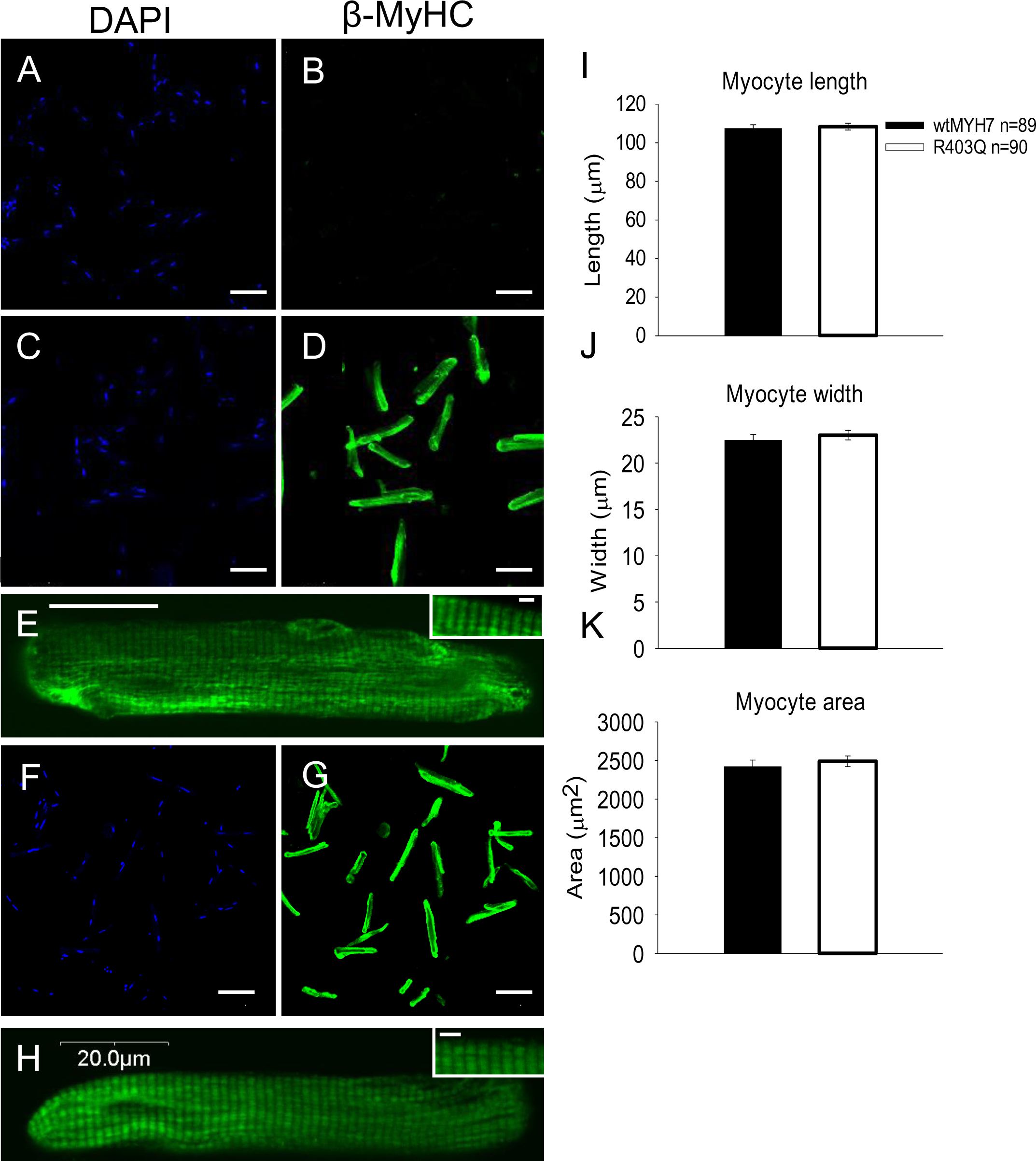
Highly efficient gene transfer of the normal and mutated human β-MyHC protein did not affect sarcomere structure or morphology. Myocytes were fixed on day 3 in culture and the nuclei were stained with DAPI and β-MyHC protein expression was probed with a specific antibody. **(A&B)** Non transduced control rat myocytes do not express any detectable level of β-MyHC protein. Scale bar=100μm. **(C&D)** On day 3 in primary culture wtMYH7 expression was apparent in >98% of the myocytes. Scale bar=100μm. **(E)** Normal wtMYH7 β-MyHC expression was homogeneous across the length and width of the cardiac myocyte. Scale bar=20μm. Inset scale bar=2.0μm **(F&G)** R403Q mutant myosin gene transfer was equally efficient. Scale bar=100 μm. **(H)** The R403Q mutant myosin incorporated homogeneously across the length and width of cardiac myocytes just as the wtMYH7. Inset scale bar=2.0μm. **(I-K)** Myocyte length, width and area were similar in each group. Data are mean +/− SEM.

#### Myocyte morphology

Coverslips of live myocytes were visualized on an inverted microscope (10X) and images were captured using a monochromatic digital camera. The system was calibrated, and myocyte dimensions were analyzed with Image Pro Express software (Media Cybernetics, Silver Spring, MD).

#### Statistical analysis

Results are expressed as mean ± SEM. For comparison between wtMYH7 and R403Q transduced myocytes two-tailed independent *t* tests were used (*P<0.05). Significant differences are denoted by * in the figures.

## Results

### Protein expression and cell morphology

Efficient *in vitro* gene transfer of the human β-MyHC molecule is summarized in Figures 1 and 2. Here, vector-based gene transfer was highly efficient with > 98% of the adult cardiac myocytes transduced. Western blotting in Figure 1 demonstrates that β-MyHC protein expression increased over time in culture following gene transfer and was maximal at day 3. This was the case for both the wtMYH7 and the R403Q mutant myosin. Importantly, total myosin content did not increase over time in culture, indicating that the endogenous sarcomeric α-MyHC motor protein was replaced stoichiometrically in the sarcomere by the Ad5-mediated transgene product. The stoichiometric replacement of the endogenous α-MyHC with human β-MyHC was estimated at ∼40-50% in the wtMYH7 and the R403Q mutant myosin transduced myocytes by Day 4 after gene transduction (supplemental figure 1). This is in close agreement with previous work using human β-MyHC gene transfer ^31^. In addition, agreement with extensive previous works ^8,25,36–40^, gene transfer to cardiac myocytes in primary culture had no detachable effects on the expression of other key myofilament proteins (Figure 1B). Thus, in this experimental context, a direct comparison can be made of the specific effects between human wtMYH7 and the R403Q mutant myosin motors incorporated into intact adult cardiac myocytes.

Consistent with previous data, no β-MyHC was detected in non-transduced control rat myocytes (Figure 2, panel B) ^31^, and human β-MyHC protein was found to be expressed in >98% of myocytes transduced with the wtMYH7 or R403Q viruses (Figure 2, panels D&G). β-MyHC expression in each case was homogenous along the length and width of myocytes (Figure 2, panels E&H). The R403Q mutant myosin incorporated into the adult cardiac sarcomere just as the wtMYH7 molecule and did not have any detected effect on sarcomere structure or myocyte morphology (Figure 2H). Myocyte dimensions and areas were not different between the groups (Figure 2, panels I-K).

### R403Q mutant myosin effects on cardiac myofilament tension development

Next, we determined the effect of human R403Q mutant β-MyHC expression on cardiac myofilament function. Permeabilized myocytes were used to directly measure myofilament and myosin function under steady state isometric conditions. Calcium sensitivity of isometric tension development, maximal tension and isometric cross-bridge cycling kinetics were determined. Myofilament calcium sensitivity was elevated by R403Q myosin expression compared to wtMYH7 as demonstrated by a leftward shift of the sigmoidal tension-pCa relationship (Figure 3A&B). Maximal calcium activated tension, however, was significantly depressed in R403Q myocytes (Figure 3C). In calcium-activated permeabilized myocytes, the rate constant of tension redevelopment (k_tr_) was determined. Kinetically, R403Q expression increased maximal calcium-activated isometric cross-bridge cycling kinetics (Figure 4). Cross-bridge cycling was also faster at sub-maximal levels of calcium activation in myocytes expressing the R403Q mutant myosin, as compared to myocytes expressing wtMYH7 (Figure 4B).

**Figure 3.**
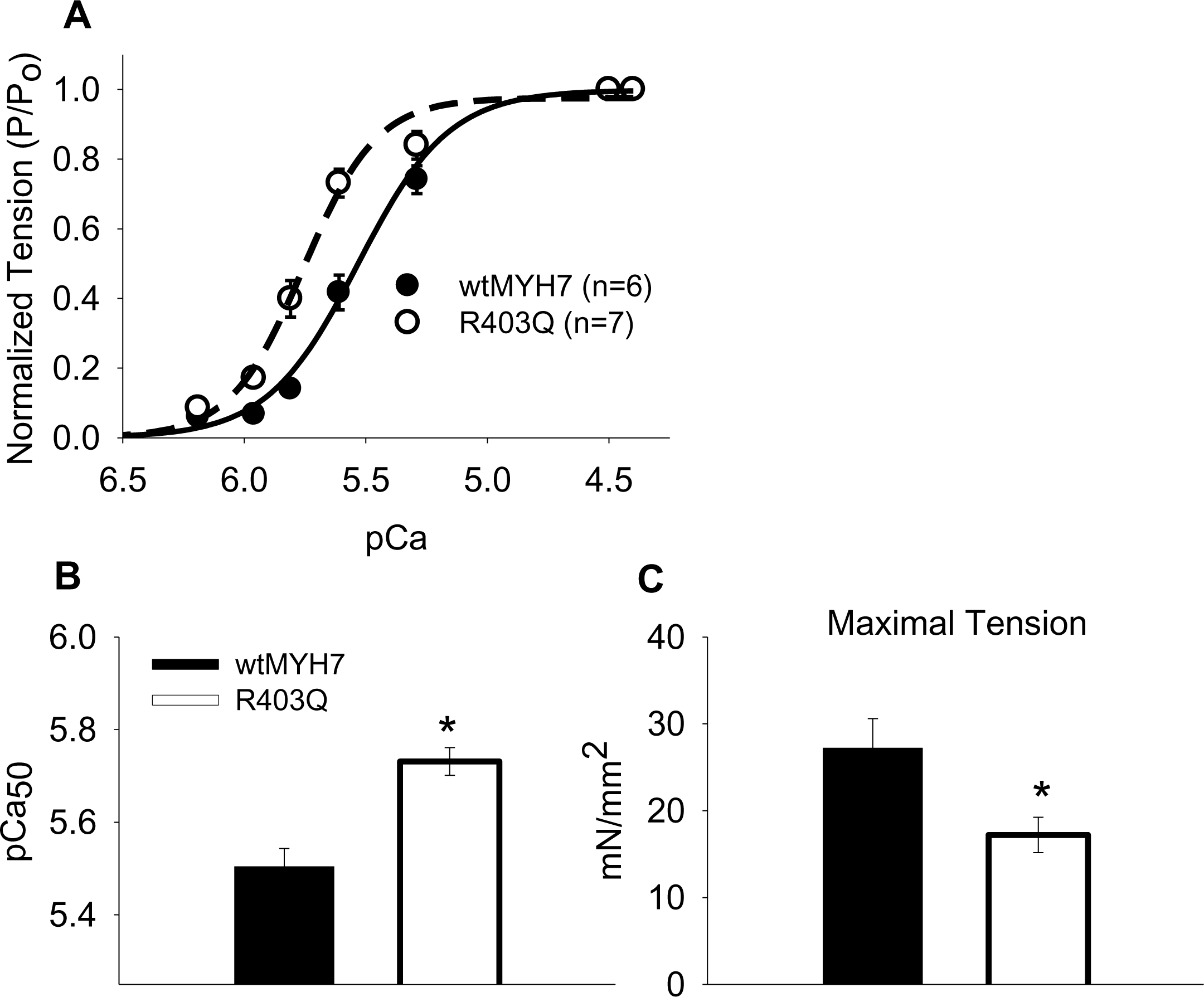
Myofilament calcium responsiveness in wtMYH7 and R403Q transduced myocytes. **(A)** Summary tension-pCa curves for each experimental group show a significant leftward shift of this relationship in R403Q myocytes. This indicates enhanced calcium sensitivity of tension in R403Q myocytes. **(B)** Significantly less calcium was required to half maximally activate tension (pCa_50_) in R403Q myocytes (pCa_50_=5.73±0.03 vs. 5.50±0.04). **(C)** Maximal calcium activated (pCa 4.5) tension was lower in R403Q myocytes (17.2±2.0, n=9 vs. 27.2±3.4, n=7mN/mm^2^). Data are mean +/− SEM. * = P < 0. 05.

**Figure 4.**
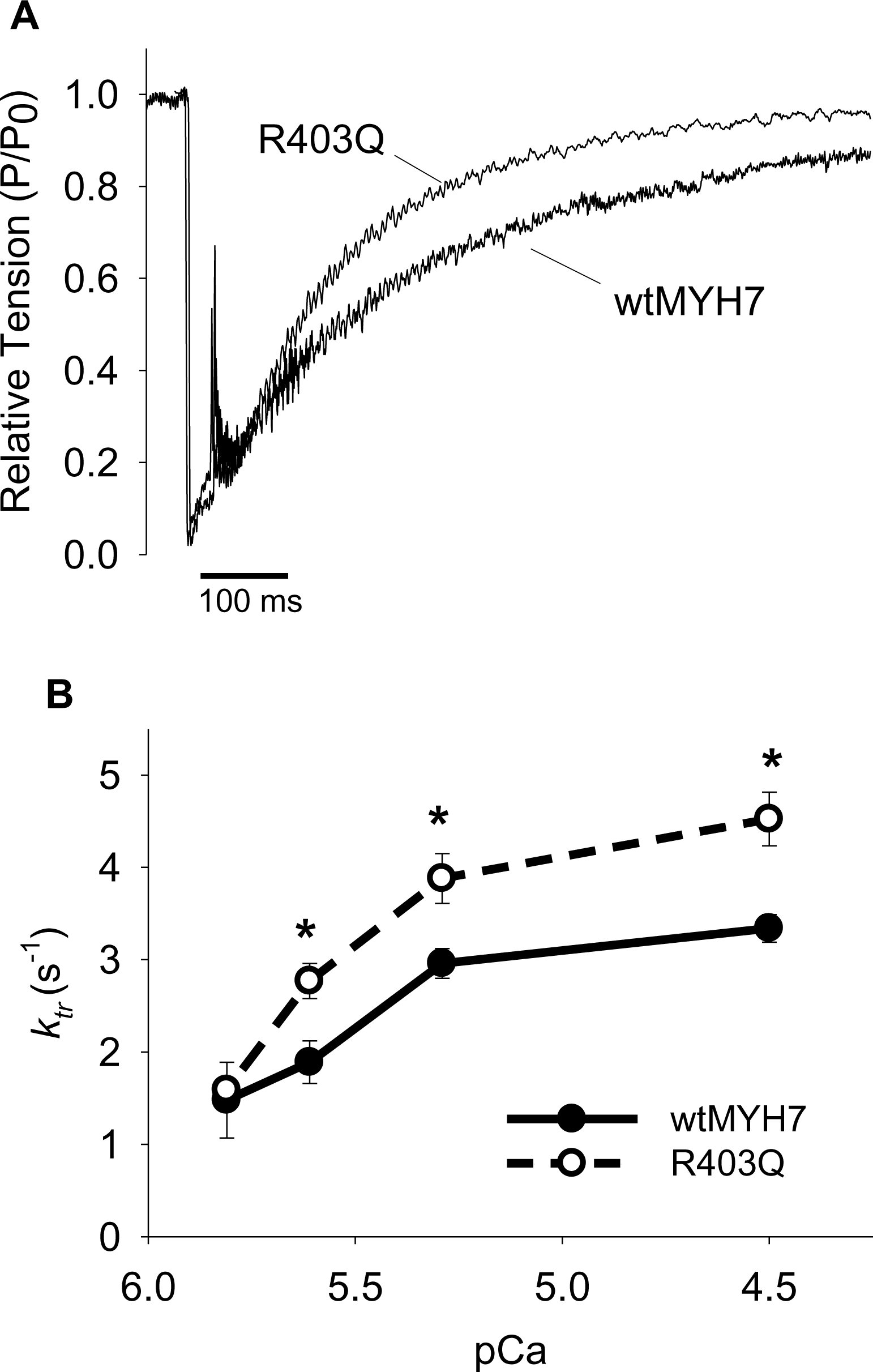
R403Q mutant myosin increases isometric cross-bridge cycling kinetics. **(A)** Original traces of maximal calcium activated (pCa 4.5) *k*_tr_ measurement in single cardiac myocytes on day 3 after gene transfer. *k*_tr_ was measured as described previously ^31,63^. **(B)** Cross-bridge cycling was faster in R403Q myocytes over a range of sub-maximal calcium concentrations. Maximal rate of cross-bridge cycling was faster (4.52±0.29s^-1^, n=9 vs. 3.33±0.15s^-1^, n=7) in R403Q myocytes. Sub-maximal calcium activated (pCa 5.3) cycling was also faster (3.88±0.27s^-1^, n=9 vs. 2.96±0.16s^-1^, n=7) in R403Q myocytes. Similarly, cycling was faster at pCa 5.6 (2.77±0.19s^-1^, n=9 vs. 1.89±0.23s^-1^, n=7). Data are mean +/− SEM. * = P <0. 05.

### R403Q mutant myosin effects on intact cardiac myocyte function

The acute effect of the R403Q mutant myosin on adult cardiac myocyte function was next determined in membrane intact cardiac myocytes stimulated electrically at 1Hz. In these experiments, physiological excitation-contraction coupling (E-C coupling) mechanisms are intact. These experiments were performed on day 3 after gene transfer when wtMYH7 or R403Q myosin expressions were maximal. Results obtained from myocytes expressing the R403Q mutant myosin were compared to myocytes transduced with the wtMYH7 adenoviral vector. R403Q myocytes were hyper-contractile and exhibited elevated calcium concentrations (Figure 5, panels A-F). Using the ratiometric calcium indicator Fura 2AM, results showed that diastolic calcium levels were higher in myocytes expressing the R403Q mutant myosin (0.78±0.009 ratio units vs. 0.61±0.01 ratio units, P<0.05). In addition, peak systolic calcium levels were higher in R403Q transduced myocytes (1.16±0.02 vs. 0.86±0.02 ratio units, P<0.05) (Figure 5). Kinetically, myocytes expressing the R403Q mutant myosin demonstrated a slower decay of intracellular calcium and sarcomere re-lengthening as evidenced by longer transient half widths (Figure 5, panels G-H). A transient half width was calculated by subtracting the time to 50% return to baseline from the time to 50% of the peak of a transient and this was shown to be significantly prolonged in R403Q myocytes (Figure 5H).

**Figure 5.**
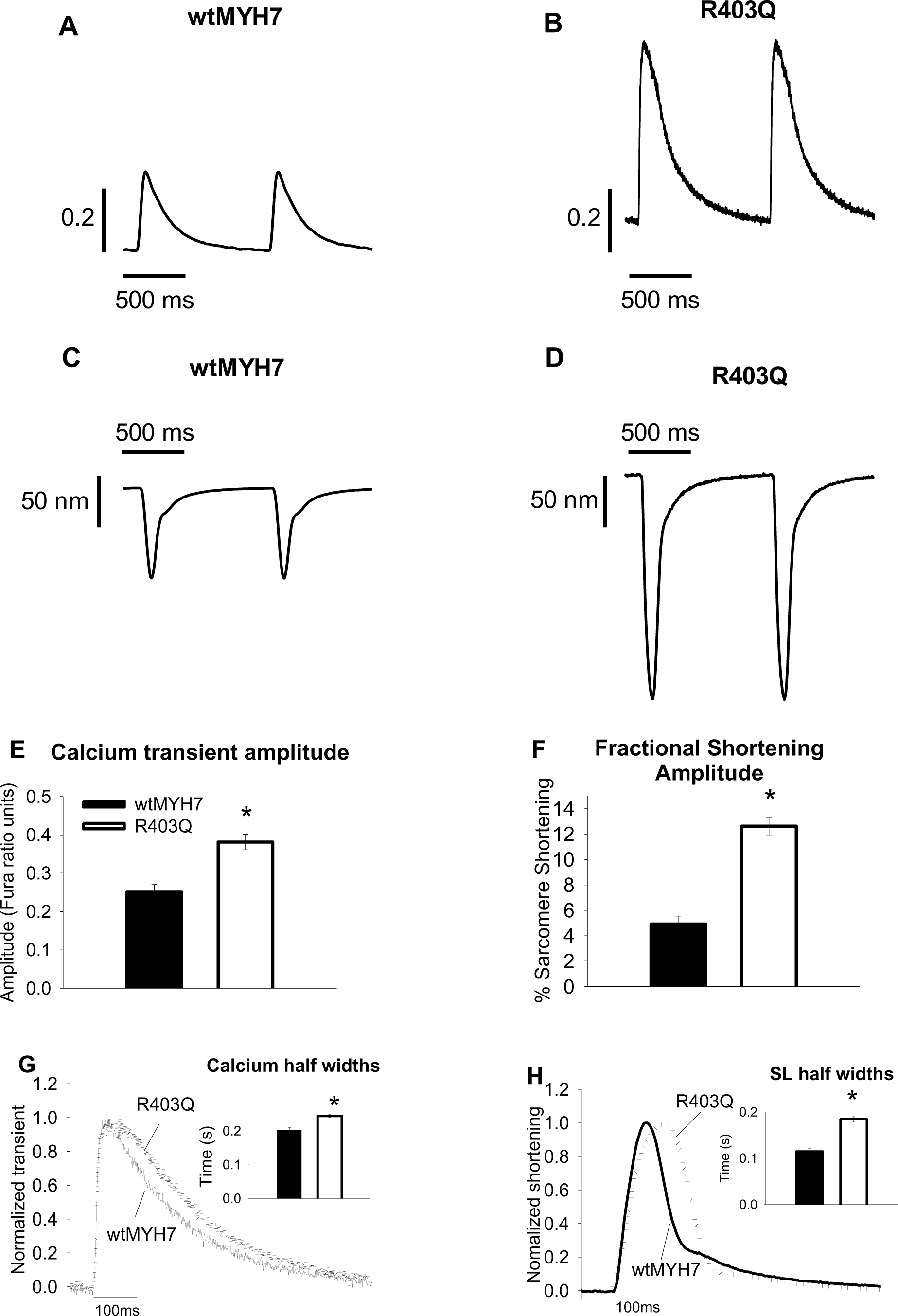
R403Q effects on intact cardiac myocyte calcium homeostasis and contractility. All data are analyzed from myocytes paced at 1Hz. **(A-D)** Original recordings of intracellular calcium transients are in A&B. The Y-axis in A and B is for the Fura ratio. Original recordings of sarcomere length shortening and re-lengthening are in C&D. **(E)** Summary of calcium transient data shows that the calcium transient amplitude was higher in R403Q myocytes (0.38±0.20, n=69 vs. 0.25±0.02, n=26). **(F)** Fractional shortening amplitude was also elevated in R403Q myocytes (12.6±0.7%, n=53 vs. 4.92±0.62%, n=26). **(G)** Normalized calcium traces show that the decay of the calcium transient was slower in R403Q myocytes (half width=0.244±0.004 s, n=69 vs. 0.201±0.008s, n=26). **(H)** Normalized sarcomere traces show that re-lengthening was also slower in R403Q myocytes (half width=0.184±0.006s, n=53 vs. 0.115±0.006s, n=26). Data are mean +/− SEM. * = P < 0.05.

The effect of R403Q mutant myosin to increase the amplitude of the intracellular calcium transient suggested that calcium release from internal stores of the sarcoplasmic reticulum (SR) may have been altered. To determine if intracellular calcium stores of the SR were different between the two groups, we used caffeine to directly measure SR calcium content in these myocytes. Consistent with the elevation of intracellular calcium levels during electrical pacing, caffeine induced release of calcium from the SR was significantly increased in R403Q transduced myocytes (Figure 6). We also determined the fractional release of calcium from the SR during electrical pacing by dividing the calcium transient amplitude during pacing by the total calcium released after the rapid application of caffeine. R403Q transduced myocytes mobilized a significantly higher fraction of releasable calcium during pacing (88.9±2%, n=8 vs. 72.1±3%, n=9)(Figure 6D).

**Figure 6.**
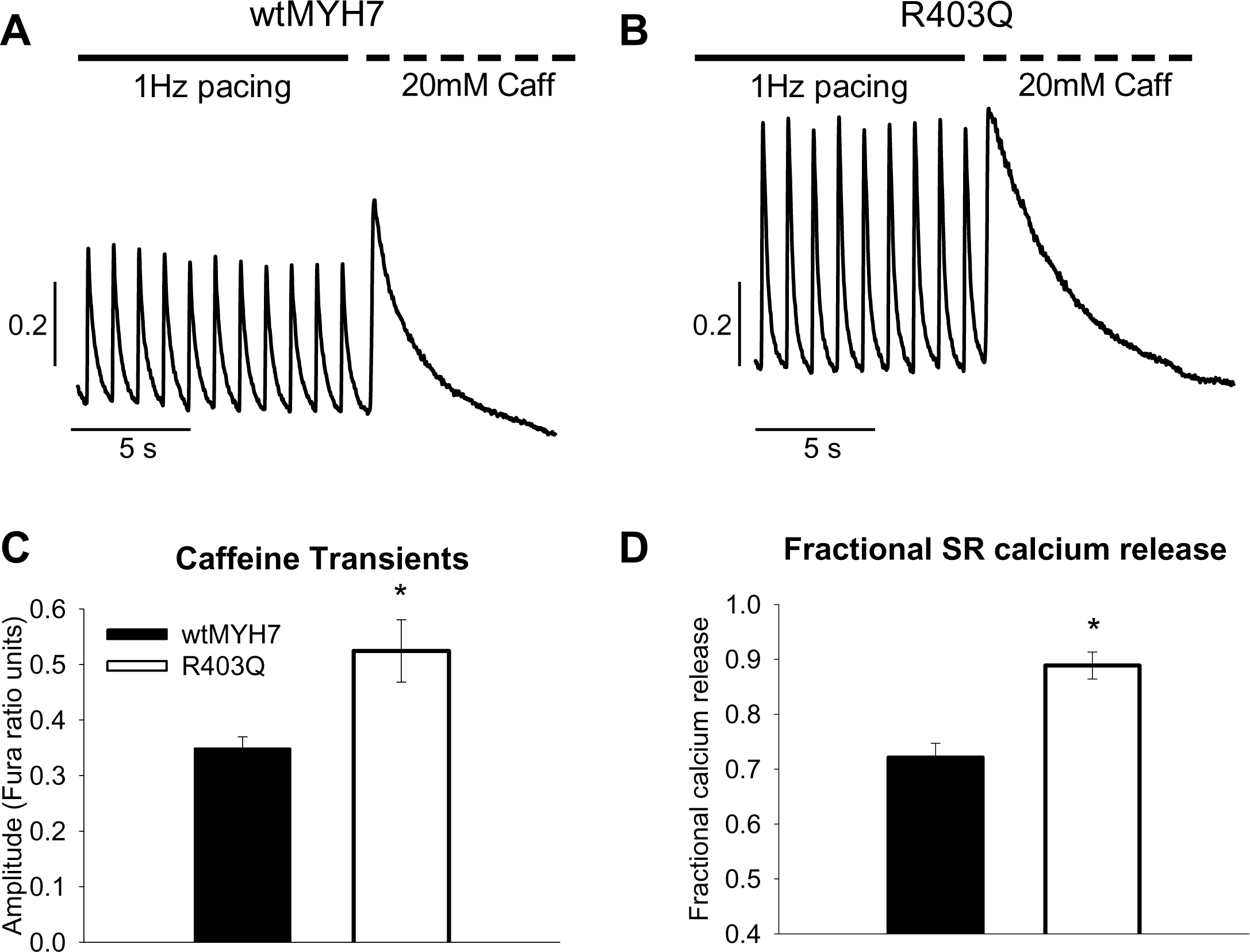
**Functional SR calcium load via rapid** caffeine application in wtMYH7 and R403Q transduced myocytes. **(A&B)** Original calcium recordings during 1 Hz pacing and with rapid application of 20mol/L caffeine for wtMYH7 and R403Q myocytes respectively. **(C)** The amplitude of calcium release by caffeine was greater in R403Q myocytes (0.52±0.06, n=17 vs. 0.35±0.02 ratio units, n=21). **(D)** Fractional calcium release during 1 Hz pacing was greater in R403Q myocytes. Data are mean +/− SEM. * = P < 0.05.

### R403Q mutant myosin effects on myocyte β-adrenergic responsiveness

HCM is the most common cause of exercise-mediated sudden cardiac death in young competitive athletes ^3^. Therefore, we examined β-adrenergic responsiveness in myocytes transduced with the HCM causing R403Q mutant myosin. Intracellular calcium and sarcomere length shortening were recorded (1Hz pacing) before and after treatment with isoproterenol (10nmol/L). Myocytes transduced with wt*MYH7* responded normally to isoproterenol treatment and calcium transients and contractility were both increased after β-adrenergic stimulation (Figure 7, panels A-D). On the other hand, R403Q transduced myocytes did not respond normally to isoproterenol. Following isoproterenol treatment R403Q myocytes displayed instability in calcium homeostasis as evidenced by significant calcium release events between electrical stimuli (Figure 7, panels E-H). The release of calcium between electrical stimuli (arrows in Figure 7, panel G&H) was significant enough to also cause spontaneous contractions. The improper release of calcium between electrical stimuli resulted in an impaired response to isoproterenol treatment (Figure 7I-J). Neither calcium transient amplitude nor contraction amplitude were increased following β-adrenergic stimulation in R403Q transduced myocytes (Figure 7I-J).

**Figure 7.**
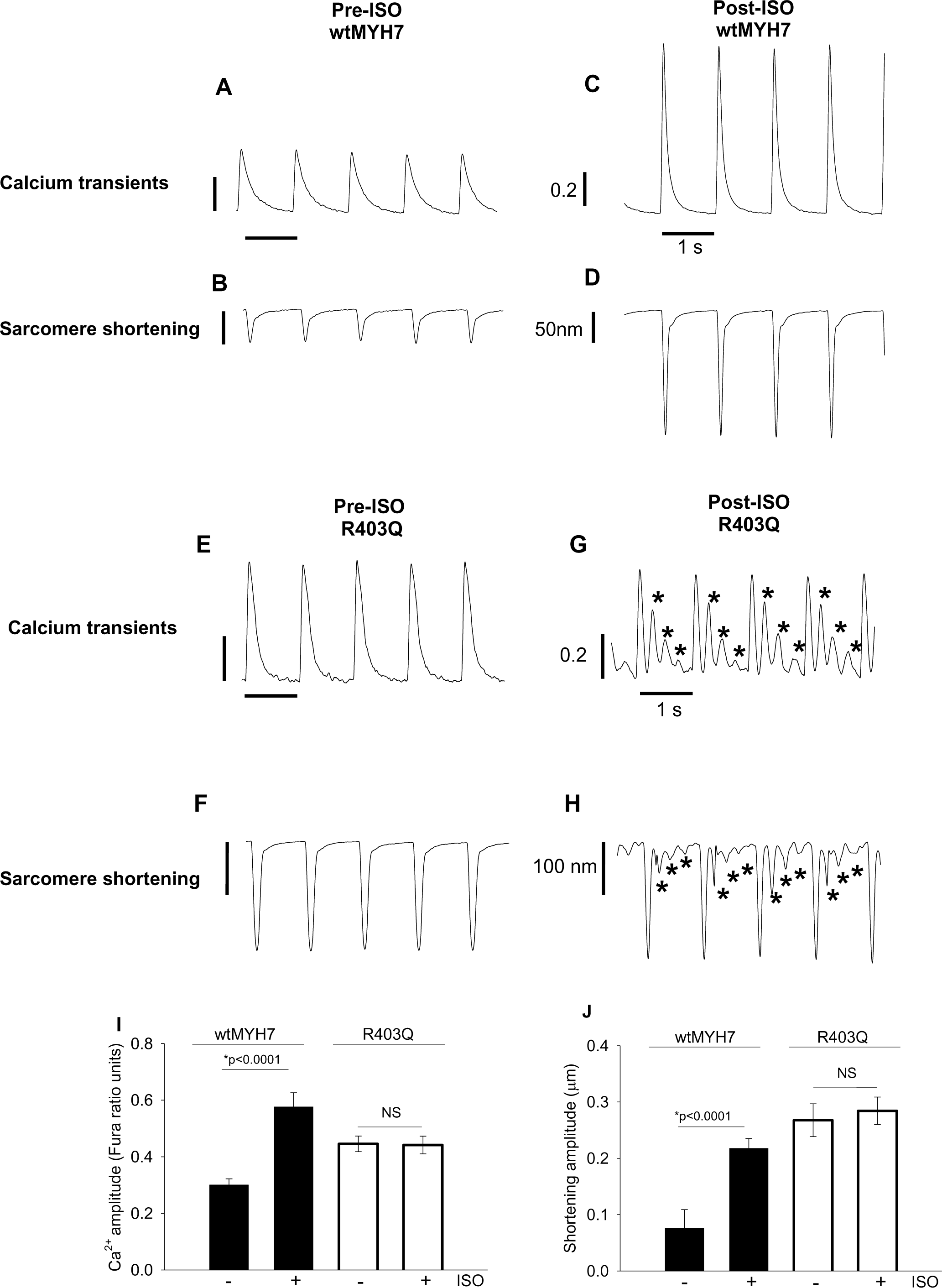
β-adrenergic responsiveness in wtMYH7 and R403Q transduced myocytes. **(A-D)** Top panel shows original calcium and sarcomere shortening traces from wtMYH7 transduced myocytes before and after treatment with isoproterenol (10nmol/L). **(E-H)** Bottom panel shows original calcium and sarcomere shortening traces from R403Q transduced myocytes before and after isoproterenol treatment. wtMYH7 transduced myocytes responded normally to isoproterenol**. Panel G** shows that R403Q myocytes generated spontaneous calcium transients (asterick, *) in between electrical stimuli. **Panel H** shows that contractions also occurred in between electrical stimuli due to unstable calcium homeostasis. **(I)** In wtMYH7 myocytes calcium transient amplitude (1hz pacing) was elevated after isoproterenol treatment (0.30±0.02 ratio units vs. 0.58±0.05, n=28). In R403Q transduced myocytes, however, calcium transient amplitude was not greater after isoproterenol treatment (0.45±0.03 vs. 0.44±0.03, n=17). **(B)** In wtMYH7 transduced myocytes the amplitude of contraction increased from 71.0±20.0 nm to 219.0±8.0 nm after isoproterenol stimulation. On the other hand, R403Q myocytes did not experience augmentation of contractility after isoproterenol treatment (253±20.0 nm vs. 219.0±20.0 nm). Data are mean +/− SEM. * = P < 0.05.

### Spontaneous calcium release (SCR) after a change of pacing frequency

Next we tested the response of R403Q myocytes to a rapid change in the pacing protocol as done before to probe arrhythmia mechanisms using Purkinje cardiac cells ^41^. Myocytes were paced to 3Hz and then were immediately slowed to 0.5Hz. Typical original recordings are shown in Figure 8. Spontaneous calcium release (SCR) and contractions were observed in ∼65% of R403Q myocytes and in less than 1% of wtMYH7 transduced myocytes.

**Figure 8.**
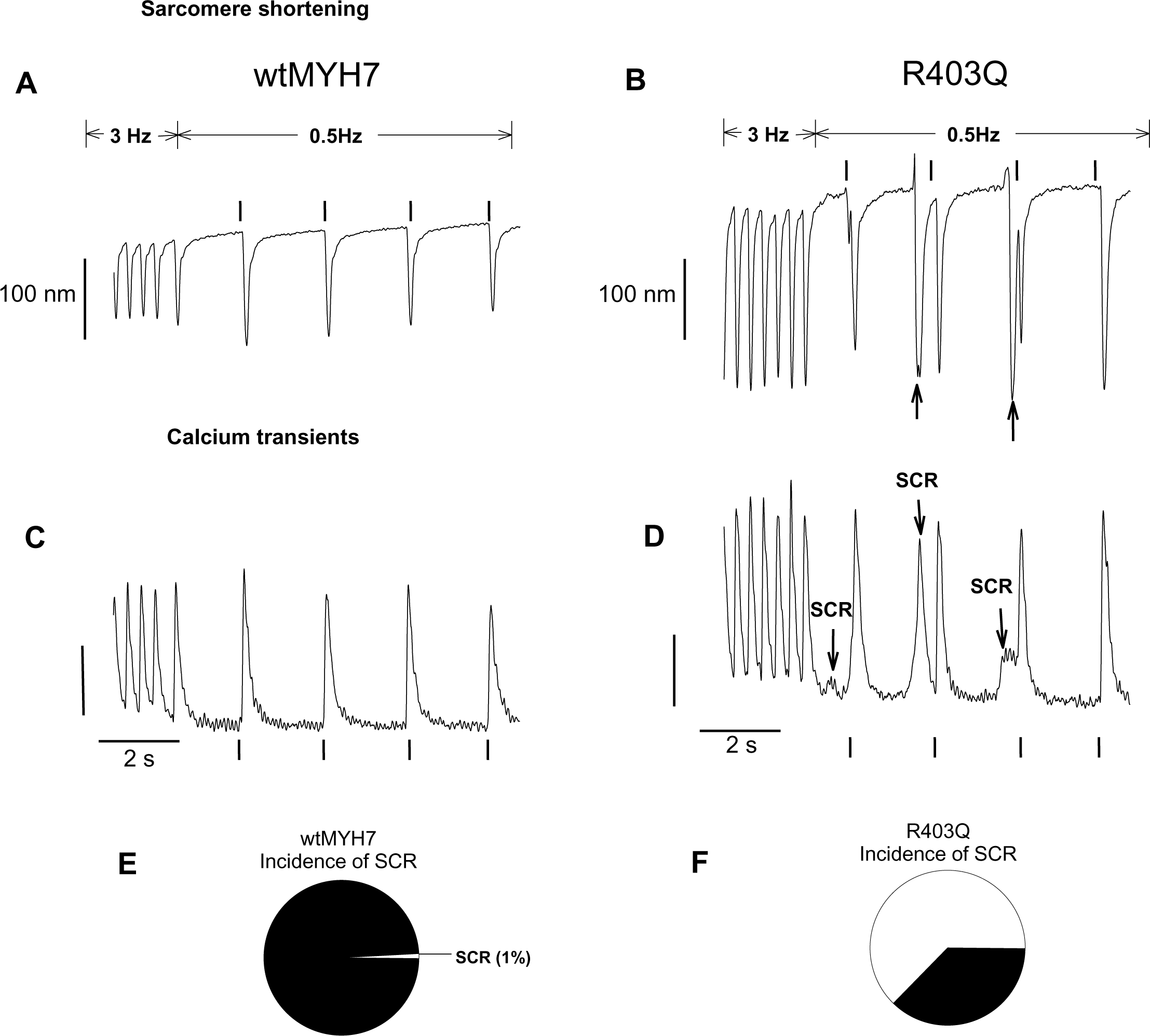
Greater incidence of spontaneous calcium release (SCR) and contractions in R403Q myocytes by rapid slowing of pacing frequency. **(A&B),** Original sarcomere shortening recordings in myocytes paced at 3Hz and then switched to0.5 Hz. Spontaneous contractions were frequently observed in the R403Q myocytes (B), but not in the wtMYH7 transduced myocytes (A). **(C&D),** simultaneous calcium transient measurement in wtMYH7(C) and R403Q myocytes(D). Spontaneous calcium release events commonly occurred in R403Q myocytes (denoted by arrows). wtMYH7 myocytes rarely showed SCR when pacing was decreased from 3 to 0.5Hz. **(E),** Less than 1% of wtMYH7 myocytes (1/15) demonstrated SCR and spontaneous contractions. **(F),** 10/16 or 65% of R403Q myocytes demonstrated SCR and spontaneous contractions.

## Discussion

Here we used acute gene transfer to study the direct functional effects of the cardiomyopathy-causing human R403Q mutant β-MyHC in otherwise normal adult cardiac myocytes. Results demonstrate that the human R403Q mutant myosin readily incorporates in the cardiac sarcomere, with ∼40% of the endogenous sarcomeric myosin stoichiometrically replaced. This extent of stoichiometric replacement was like gene transfer of normal human β-MyHC reported before.^31^ Human R403Q mutant β-MyHC did not directly cause sarcomere disarray or cellular hypertrophy in primary culture conditions. Notably, in electrically stimulated intact adult cardiac myocytes, the transduced myocytes were hyper-contractile and intracellular calcium concentration increased due to incorporation of human R403Q mutant motor. At the sub-cellular myofilament level in permeabilized myocytes, the R403Q mutation increased calcium sensitivity and significantly accelerated cross-bridge cycling kinetics. These results indicate that a disease-causing alteration of cardiac myosin structure affects myofilament calcium responsiveness and can have significant effects on intracellular calcium handling, especially following β-adrenergic stimulation. Increases in intracellular calcium concentrations may provide a trigger for the ensuing cardiac hypertrophy, cellular disarray and susceptibility to arrhythmia that is characteristic of the human HCM phenotype.

### R403Q effects on myocyte morphology and sarcomere structure

Under primary culture conditions, wherein the adult myocytes are stable and quiescent, data show the mutant R403Q myosin incorporates into the sarcomere normally and in a similar manner to wtMYH7. Thus, the characteristic HCM disease phenotype of sarcomere and myocyte disarray ^42,43^ are likely a secondary compensatory mechanism to account for the mutant myosin molecule expression. This is consistent with other work showing that the R403Q mutant myosin expressed in rat ventricular cardiac myocytes resulted in normal thick filament assembly, and produced myofibrils with well-defined I bands, A bands and H zones ^44^. On the other hand, in COS cells ^45^ and in feline myocytes ^46^ there was some evidence of myofibril and sarcomere disarray caused by expression of the human R403Q mutant myosin. In the case of the feline myocyte study, however, it is important to point out that sarcomere structure was normal 48 hours after gene transfer and that sarcomere disarray was only apparent 120 hours after mutant myosin gene delivery and expression. Thus, the work from Marian et al. ^46^ is consistent with the hypothesis that hypertrophy and sarcomere disarray characteristic of HCM may be secondary and compensatory to mutant myosin expression. Alternatively, the effect of the mutant myosin molecule on sarcomere structure and cellular organization may be more apparent when the mutation is expressed during cardiogenesis. Support for this idea has been reported in a zebrafish model of hypertrophic cardiomyopathy caused by a cardiac troponin T (tnnt2) gene mutation ^47^. Becker et al., ^47^ reported that sarcomere mutations can alter cardiac myocyte structure and function at the earliest stages of heart development with distinct effects from adult hearts.

In this setting, using genetic engineering platforms, sarcomere mutations in HCM patient specific induced pluripotent stem cell derived cardiac myocytes (hiPSC-CMs) may provide for deeper mechanistic insight ^48–52^. In this light, several studies have been conducted on HCM patient generated hiPSC-CMs. One key advantage of this system is the beating hiPSC-CM are comprised of human genes expressing human proteins organized into human sarcomeres. Here, the hiPSC-CMs express the β-MyHC gene, which is the dominant motor protein isoform expressed in the adult human heart. HCM mutations in human β-MyHC, when compared to isogenic lines without the mutant, shows evidence of hyper-contractility and discordant Ca^2+^ signaling resulting in cellular arrhythmias ^49^. Although these hiPSC-CM HCM studies can provide mechanistic insights, they also demonstrate limitations of this system including phenotypic variability between lines that typically necessitate individual functional assessments. Nonetheless, these hiPSC-CM studies do offer an exciting platform, especially to combine with animal model system as done here, to make mechanistic insights into the cellular and molecular etiology of HCM.

### R403Q effects on myofilament function and myosin cross-bridge cycling

We determined isometric force production, calcium sensitivity of tension and the kinetics of acto-myosin cross-bridge turnover in permeabilized (skinned) cardiac myocytes. Cardiac myocytes were transduced with either wtMYH7 or the HCM causing R403Q mutant myosin gene to assess myosin and myofilament function more directly. The results provide evidence that the R403Q mutant myosin affects the transition of acto-myosin cross-bridges between non-force generating and force generating states. Here we found that the R403Q mutant myosin accelerated isometric cross-bridge cycling at both maximal (pCa 4.5) and sub maximal levels of calcium activation. According to a simple 2-state model of acto-myosin interactions ^53^ in which the formation of force-generating cross-bridges occurs via an apparent rate constant *f*_app_, and cross-bridge detachment occurs with apparent rate constant *g*_app_, the overall cross-bridge cycling rate is given by *f*_app_ + *g*_app_. Therefore, in this formalism, R403Q increases cross-bridge cycling by increasing cross-bridge attachment (*f*_app_) and/or detachment (*g*_app_).

More insight into whether *f*_app_ or *g*_app_ are affected by R403Q can be gained upon examination of total isometric tension development. In the simple 2-state model, total isometric tension is proportional to *f*_app_/(*f*_app_ + *g*_app_). At sub maximal calcium activation levels, it is likely that *f*_app_ is increased to a greater extent than *g*_app_ as indicated by greater total tension development and faster cross-bridge cycling. On the other hand, under conditions of maximal calcium activation (pCa 4.5), it is likely that *g*_app_ is increased to a greater extent as indicated by a reduction in total tension development and faster cross-bridge cycling. Consistent with our data under maximal calcium activation conditions, a study that utilized single cardiac myofibrils from HCM patients with the R403Q genotype reported that maximal calcium activated cross-bridge turnover was accelerated and maximal tension was depressed ^54^. This apparent calcium dependent effect of the R403Q mutant myosin on myofilament function is consistent with a previous study that used skinned papillary muscle strips from the R403Q transgenic mouse model ^15^. Blanchard et al ^15^ reported that at sub maximal calcium activation isometric tension was ∼3 times higher in mutant than in wild-type muscle strips. Maximal calcium activated tension was, however, marginally lower. Thus, in the case of the R403Q mutant myosin heads may accumulate in a state that promotes activation of the thin filament at sub maximal calcium concentrations but lowers maximal tension when the thin filament is fully activated by calcium.

Our results can be further discussed in the context of recent work providing evidence of a new layer of myofilament-based regulatory complexity, and is pertinent to understanding effects of mutant myosin motors on cardiac contraction ^55–57^. Data suggest a significant fraction of myosin heads are in a sequestered state with an extremely low ATPase rate and not capable of interacting with actin. This state has been referred to as the super relaxed state of myosin (SRX). In comparison, the disordered state of myosin (DRX) has a higher ATPase rate and has the capacity to transition to actin interacting cross-bridge states. In this framework, recent hiPSC-CM studies provide evidence that the R403Q mutant myosin serves to shift myosin states from the SRX to the DRX states ^17^. This effect of mutant R403Q myosin could therefore provide the molecular mechanism underlying our findings of increased weak to strong cross-bridge state transition (k_tr_) in adult cardiac myocytes incorporating HCM mutant R403Q myosin.

The myosin R403Q mutation results in the substitution of a highly conserved neutral glutamine residue with a positively charged arginine residue in the actin binding region of the myosin head ^58,59^. This charge change in the functionally critical actin binding domain may promote force generation when the thin filament is only partially activated by interactions with one or more thin filament proteins. This potential interaction may dissipate when the thin filament is maximally activated by calcium, thereby decreasing the number of strongly bound cross-bridges during maximal calcium activation. It is important to point out that cardiac myofilaments normally operate at sub maximal levels of calcium activation when activated electrically by physiological excitation contraction coupling mechanisms.

### R403Q effects on intact myocyte contractility and calcium homeostasis

Effects of R403Q mutant myosin on myosin and myofilament function were also observed in the more physiologically relevant context of the membrane intact adult cardiac myocyte. Consistent with the hyper-contractile phenotype in skinned myocytes under sub maximal calcium activating conditions, R403Q myocytes were also hyper-contractile when activated electrically. The hyper-contractility observed in membrane intact myocytes may be partially due to increased myofilament calcium responsiveness and accelerated cross-bridge turnover, as discussed above. These primary alterations of myofilament function apparently also affected intracellular calcium homeostasis and the observed elevations of intracellular calcium concentrations could account for most of the hyper-contractile phenotype. The mechanism contributing to elevated calcium levels is unclear but may involve the R403Q mutant effects on myofilament calcium responsiveness and thin filament activation. For example, faster and greater extent of thin filament activation by R403Q myosin cross-bridges may alter the kinetics and extent of calcium release from the SR. Caffeine experiments performed here suggest that this may be the case. Data show that the amount of calcium released from the SR, as well as the fractional release of calcium during electrical pacing were each elevated in R403Q myocytes. Therefore, myosin R403Q transduced myocytes are subject to calcium overload and are at risk of becoming arrhythmic and switching on calcium sensitive hypertrophic signaling pathways.

The mechanism by which a mutation in the myosin motor protein could alter SR calcium handling function is not known. Because the experimental design of myocytes transduced in primary culture it would seem the impetus for secondary compensatory adaptions would be rather minimal, as we have shown previously using other mutant sarcomeric proteins studied under these same conditions ^60,61^. This contrasts with murine transgenic approaches in which literally billions of cardiac contractile cycles are present during the course of the experimental timeline, and this clearly would be a major substrate for adaptations. One could speculate that the heightened myofilament calcium sensitivity shown here for myosin R403Q could serve as a calcium trap that, in turn, could influence calcium handling, as proposed by others ^23^.

Because HCM is the most common cause of sudden cardiac death in competitive athletes during athletic performance, we performed experiments to test β-adrenergic responsiveness in R403Q myocytes. These experiments mimic the clinical stress test that is used by cardiologists to reveal cardiac arrhythmias and cardiomyopathy. Intracellular calcium homeostasis was adversely affected following β-adrenergic stimulation. Calcium release from the SR became very unstable and calcium transients large enough to cause contractions were apparent in between electrical stimuli in R403Q myocytes. In control experiments, wtMYH7 transduced myocytes responded normally to isoproterenol treatment. On the other hand, R403Q myocytes failed to increase contractility following isoproterenol treatment. The lack of response to isoproterenol may be attributed to the high level of contractility prior to isoproterenol treatment or to the leak of calcium from the SR in between electrical impulses following isoproterenol treatment. Intracellular calcium homeostasis was also apparently altered following a rapid change in pacing frequency from 3 to 0.5Hz. Collectively, the *in vitro* stress tests indicate that the mutant myosin causes increased incidence of arrhythmias mediated by calcium mishandling.

One limitation of the present study involves using an animal experimental platform to study a human disease. Strengths and limitations of such an approach are well noted. Specifically, we used rodent myocytes in which to express and incorporate into the sarcomeres the human myosin motor protein. Here, the mutant motor was from the correct species, and it was the correct isoform (*MYH7*); however, this was then expressed in rodent adult myocytes in which the *MYH6* isoform is expressed, and not the *MYH7* isoform ^9,25,62^. To help mitigate this limitation, all comparisons were made in this work between myocytes incorporating either the human mutant R403Q in the *MYH7* backbone or, with similar incorporation amounts, via human wtMYH7. In this way, differences reported can be attributed to the human mutant R403Q protein.

In summary, direct gene transfer of normal human *MYH7* or HCM mutant R403Q *MYH7* was conducted to provide additional insights into direct effects of the human HCM mutant myosin on adult myocyte function. Data provide evidence of stoichiometric myofilament incorporation of the mutant molecule like normal β-MyHC that, in turn, caused heightened myofilament calcium sensitivity with reduced maximum tension and faster cross-bridge cycling kinetics, all independent of altered sarcomeric or cellular structure. These results are consistent with recent work showing the R403Q mutant myosin to shift myosin cross-bridge regulatory states from SRX to DRX states ^17^. The mutant myosin caused altered calcium handling and increased cellular contractile aftercontractions that were provoked by pacing and by adrenergic stimulation. These findings add to the literature demonstrating the primary defect of HCM R403Q myosin and contribute insights into how HCM mutants residing in the sarcomere could elicit altered calcium handling and arrhythmia in the human heart.

## Acknowledgements

This work was supported by the National Institutes of Health (Fellowship HL080880 to TJH and grant HL132874 to JMM) and the American Heart Association (Scientist Development Grant 0735464Z to TJH). This work also utilized the Morphology and Image Analysis Core of the Michigan Diabetes Research and Training Center funded by NIH5P60 DK20572 from the National Institute of Diabetes & Digestive & Kidney Diseases.

---

**Supplemental Figure 1.**
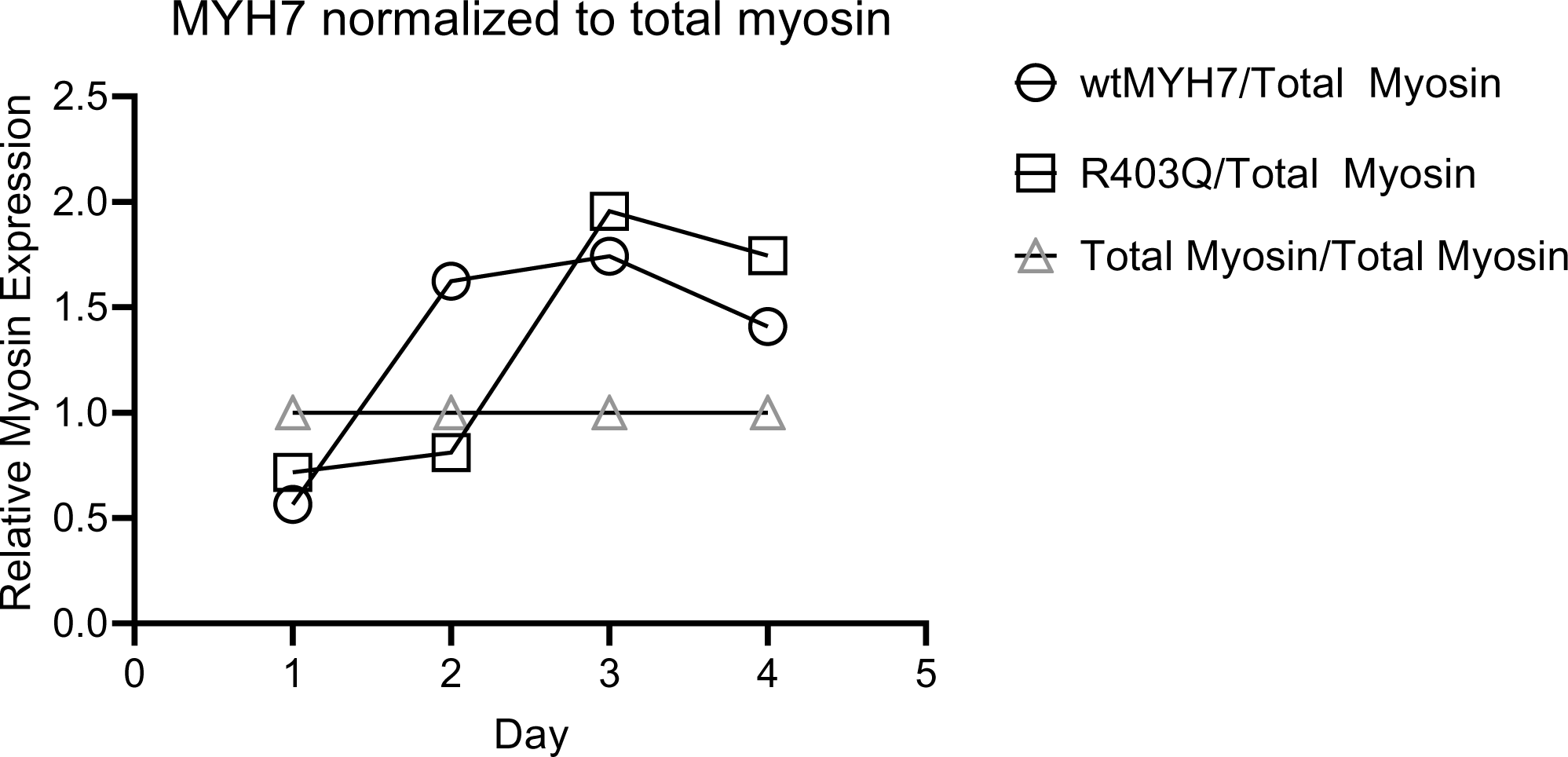

